# Detection of Static, Dynamic, and No Tactile Friction Based on Non-linear dynamics of EEG Signals: A Preliminary Study

**DOI:** 10.1101/2020.04.05.026039

**Authors:** Golnaz Baghdadi, Mahmood Amiri

## Abstract

Touching an object leads to a frictional interaction between the skin and the object. There are two kinds of friction: the first contact that leads to static friction and the dragging phase that leads to dynamic friction. No study has been performed to show the effect of friction type on EEG signals. The main goal of the current study is to investigate the effect of tactile friction on non-linear features of EEG signals.

Participants performed a tactile task that each of its trials had three states: the sensation of 1) static friction, 2) dynamic friction, and 3) no friction. During the experiment, EEG signals were recorded, and different linear and non-linear EEG indices were extracted and analyzed to find the effect of the tactile friction on EEG signals.

Linear features such as spectral features were not a good choice to distinguish between the states. However, non-linear features such as Lyapunov exponent, Higuchi’s dimension, and Hurst exponent had the potential to separate the mentioned states. Results also showed signs of predictability (negative Lyapunov exponent) in the signals recorded during dynamic friction and the existence of long-range dependency (memory) in EEG signals recorded during all states. The complexity of the tactile system in Theta band was also higher than the Delta band. The results of this research not only increase our knowledge about brain non-linear dynamics in response to tactile friction but also lead to a design of a preliminary system that can automatically detect friction between the skin and surfaces.

## 1. INTRODUCTION

Due to the collision of the skin with a surface, tactile friction is formed between the skin and the surface. The frictional interaction between the skin and the surface influences the tactile perception [1, 2]. Indeed, the level of friction or the degree of roughness affects the characteristics of this perception [3] and, consequently, the brain state [4]. The activation of somatosensory cortex, posterior cingulate cortex, central-parietal sites, lateral parietal operculum, insula, lateral prefrontal cortex, and supplementary motor area have been reported by several research during tactile roughness discrimination [5-7]. Some differences were also observed between event-related potential (ERP) components (N100, N200, and P300) during touching surfaces with different roughness levels [8]. The results of these studies have been used to design tactile brain-computer interfaces (BCI) [9, 10] systems. Alpha and Theta are two brain frequency bands that their activations have been reported in many studies on tactile roughness discrimination [11-14].

It was shown that the method of touching a surface: direct passive (Participant does not move his/her skin on the surface. The surface is applied to some part of the skin and then is pulled across the skin by someone else or by a mechatronic tool), direct active (Participant pulls his skin over the surface actively) [8, 15, 16], or indirect passive (Touching is done by a probe not the direct contact of the skin) [17] have considerable impact on tactile perception.

The speed of moving the body over a surface can also vary the characteristics of the tactile perception [5]. Even touching a surface with an open or closed eye has effects on texture perception [18]. It has been shown that not only the vision but also auditory stimuli can have a significant effect on tactile friction perception [19]. Therefore, tactile experiments need to be performed in a quiet room. The deficit of the motor system [20], the impairment of neuro-cognitive system [21], age and gender [22], the amount of contact force [23], and the state of the skin [1, 24] are the other factors that play significant role in tactile perception.

In addition, the type of friction is another important factor that its effect on brain dynamics needs to be investigated. There are two kinds of friction: static and dynamic. In static friction, two objects do not move relative to each other. In dynamic friction, at least one object moves over the other [25]. In a tactile experiment, as the skin contacts with an object, static friction is created between the skin and the object surface. This friction is static since skin and surface do not move relative to each other. As skin moves over the object surface, dynamic friction is created.

In our knowledge, no study has been performed to show the effect of friction type on EEG signals. In addition, the focus of all previous studies performed on tactile friction was on the frequency or ERP features. No study has been done on the effect of tactile perception on non-linear features of EEG signals. Hurst exponent, Higuchi’s dimension, Lyapunov exponent, or entropy are examples of non-linear features. These features give information about the system that produces EEG signals, and the importance of these features have been shown in different EEG studies [26, 27]. The main goals of this study is (1) to explore how the brain dynamics (linear and non-linear features) change in response to a sequence of events: touching a surface without moving the skin (sensation of a static friction), pulling the skin along the surface in the direct passive way (sensation of a dynamic friction), and disconnecting the skin from the surface (sensation of no friction), and (2) designing a system that can automatically detect the mentioned three states according to the EEG signals’ features

The results of this study can be useful for scientists in three fields: (1) in neuro-cognitive fields, the results increase the knowledge about the dynamics of the brain according to the tactile EEG signals; (2) in designing computer to brain or brain to brain interfaces (CBI and BBI), the results provide some possible EEG biomarkers that can be used in CBI or BBI as indices to design brain stimulation protocols; (3) in tactile industry, the results electrophysiologically reveal the effect of friction on brain wave signals that help scientists to find out more knowledge about clothing tactile comfort.

## 2. METHOD

### 2.1. Subjects

Nine healthy right-handed subjects (four females) with the age of 27±1.5 years voluntarily participated in our study. They had no history of neurological disorders, motor disorders, or skin diseases. They participated in a tactile sensation experiment, which was approved by the Iran University of Medical Sciences (#IR.IUMS.REC.1396.0294) and Kermanshah University of Medical Sciences (IR.KUMS.REC.1397.011). The participants were aware of the experimental procedure and were signed the informed consent form.

### 2.2. Experiment Procedures

As mentioned in the introduction part, several studies show the effect and importance of the roughness of a surface on EEG signals. Therefore, the roughness of a surface can affect and change the sensitivity of our designed system for the detection of the contact/drag phase. To reduce the effect of this issue, the experiment consisted of tactile stimuli with different levels of roughness (rough, semi-rough, and smooth). To deliver the tactile stimuli, a rotating plate that had three surfaces with three different roughness levels was designed. These surfaces were applied to the skin by a pseudo-random sequence to prevent learning/anticipation effects. To minimize the mentioned effect, the interstimulus interval was also set randomly. the plate, which consisted of three surfaces, had two degrees of freedom. It could move vertically to make a direct contact of the finger with the surfaces. It could also rotate around its axis that allowed the finger to be dragged across different surfaces passively. To eliminate the effect of visual acuity on the determination of roughness or the contact/drag phase, participants were requested to close their eyes. They sat on a comfortable chair in a quiet and free of noise room of the National Brain Mapping Lab (NBML) of Iran. The index fingers of the participants were fixed above the plate using a holder, which allowed the hand’s muscles to be in a relaxed state and helped to have an equal level of the pressure of touching for all participants. The equal pressure kept the contact force consistent across the participants and the trials. Fig. 1 shows one of the participants during the experiment and the situation of his hand in the setup. Fig. 2 schematically shows the setup of the experiment.

**Fig. 1.**
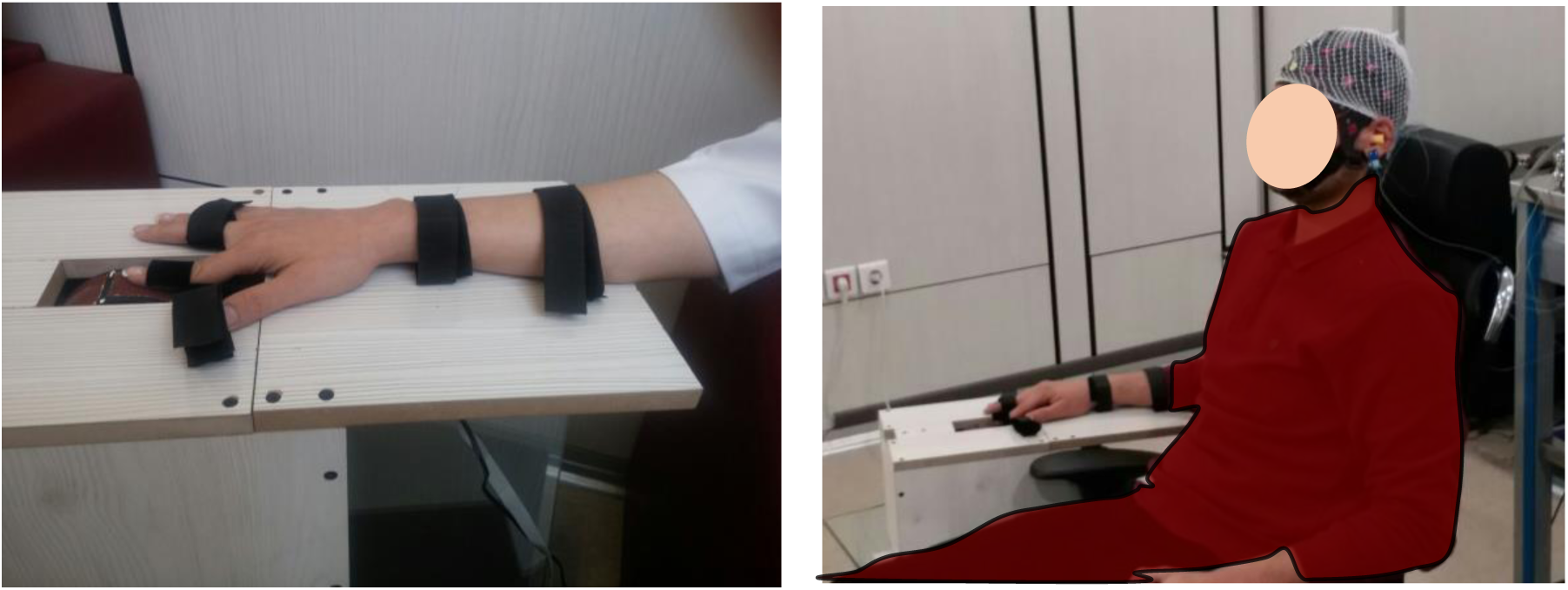
Right side: one of the participants during the experiment; Left side: the situation of his hand in the setup.

**Fig. 2.**
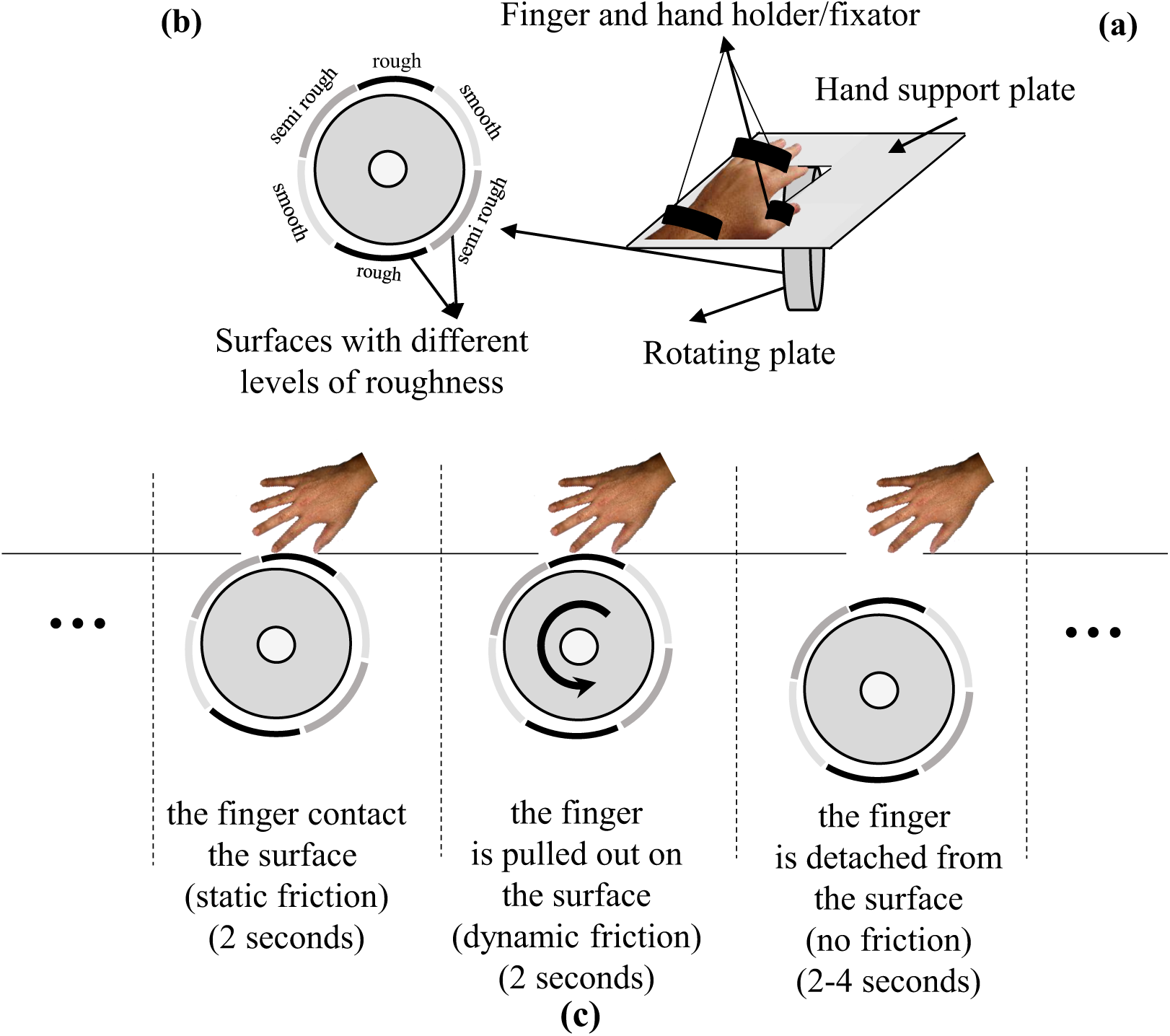
A part of the experimental setup. (a) The rotating plate and hand fixator; the plate moves and rotates below the fixed fingertip with a specific timed schedule. (b) The rotating plate with different surfaces. (c) Three blocks of each trial (contact phase (static friction), dragging phase (dynamic friction), detaching phase, (no friction)).

During the experiment, participants felt each of the three surfaces 36 times. Each trial of touching a surface can be divided into three blocks as follow:

- Block 1: In the first block of each trial, the plate moved up vertically and made direct contact with the fingertip, but did not move. Therefore, static friction was created between the fingertip and the surface for two seconds.
- Block 2: In the second block, for two seconds, the plate started to rotate, and the surface was pulled out underneath the index finger. Therefore, the tactile system perceived dynamic friction.
- Block 3: In the third block, the plate moved down vertically and thus disconnected the surface from the finger. The finger remained in this suspended state (with no friction), between two and five seconds.

The mentioned stages of the experiment were repeated for both left and right hands. In all blocks of trials, EEG signals were recorded using a 32 channels high-performance biosignal amplifier g.HIamp device based on the 10-20 system with a sampling rate of 512 Hz. The data were preprocessed by passing them from low and high pass filters, respectively, with 40 Hz and 0.5 Hz cutoff frequencies. Eye movement artifacts were rejected at the threshold level of ±70 μV.

### 2.3. System Design Procedure

As mentioned before, the current study aims to design a computerized system that can detect when an individual’s skin contacts a surface and when it is dragged across it based on the indices extracted from EEG signals. In other words, this system works based on the effects of static, dynamic, and no friction on EEG signals. One of the main steps in designing a BCI system is to find indices that can distinguish between the predefined states. We analyzed the linear and non-linear features to detect the desired states:

1. When the individual’s (right/left) index finger contacts the surface (static friction).
2. When the individual’s (right/left) index finger is dragged across the surface (dynamic friction).
3. When the individual’s (right/left) index finger is detached from the surface (no friction).

We expected that, regardless of the roughness (soft, semi-rough, and rough) of the touched surface, the designed automatic detection system could distinguish the above states with acceptable accuracy. It is also expected that the system is not sensitive to which hands are touched.

#### 2.3.1. Linear Features

Theta (4.5-8 Hz) and Alpha (8.5-16 Hz) bands are two brain waves that their significant activations have been reported in previous studies on tactile sensation and roughness discrimination [11-14]. Therefore, we extracted and examined the capability of these two bands’ energy to distinguish between the states. In addition to these two bands, we also investigated other brain waves (Delta (0.5-4 Hz) and Beta (16.5-32 Hz).

Delta, Theta, Alpha, and Beta bands’ signals of each trial, were divided into three parts based on three blocks described in the previous section (i.e., Block 1: The surface is connected to the fingertip; Block 2: The surface is moved, and the index finger is dragged across the surface; Block 3: The finger detached from the surface).

All of these stages were done for each surface (rough, semi-rough, and smooth) and each hand separately.

#### 2.3.1. Non-linear Features

In many of studies, it has been claimed that some features that are associated with the non-linear dynamics of the EEG signals are useful to represent the characteristics of these signals in different situations [27-30]. In addition to the importance of the speed and accuracy of the system response, the other challenge is reducing the number of EEG channels. This reduction has the following benefits:

- The decrement of the required calculation
- The increment of the response speed
- Reducing the required time for preparing the participant for EEG recording
- Reducing the risk of electrode failure
- Increasing the participants’ freedom of action

Therefore, in the next stage of our study, we investigated whether non-linear features can distinguish between desirable states with fewer EEG channels. We calculated the following indices from EEG signals:

##### Hurst exponent

Hurst exponent, which is calculated based on the following steps, shows the presence or absence of long-term memory or correlation in time series [30].

1. For a given time series x_n_ (n=1, 2, …, N), the mean is calculated.

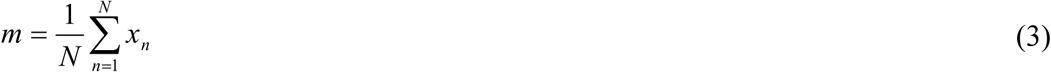
2. Each sample is subtracted from the average value calculated in the previous step.

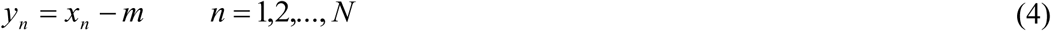
3. Then the cumulative deviate series, Z, is calculated.

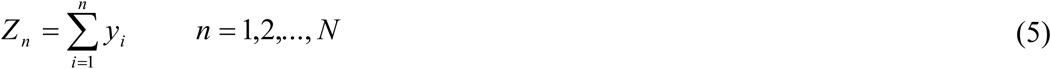
4. The range R(n) is calculated.

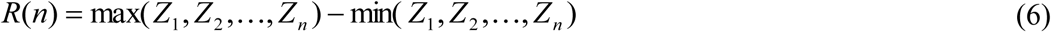
5. The standard deviation, S(n), is then calculated.

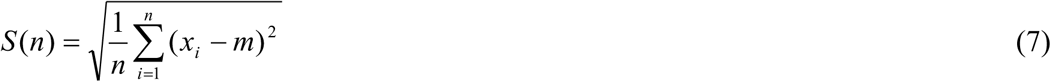
6. Calculate *R(n)/S(n)*.
7. Log[*R(n)/S(n)*] is plotted as a function of log(n).
8. Fit a straight line to the previous plotted points.
9. The slope of this line is Hurst exponent (H). In other words, fitting a power low to the expected value of *R(n)/S(n)*.

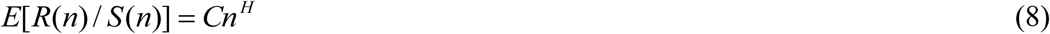

For a random time series, H=0.5. For a time series with long-range correlations H>0.5 and H<0.5 indicate the presence of long-range anticorrelation in the time series.

##### Higuchi’s dimension

Higuchi’s dimension is a measure of the complexity of a nonlinear signal. Higuchi’s dimension can be calculated according to the following steps [27]:

1. For a given time series *x*_*n*_ (*n*=1, 2, …, *N*), *k* new self-similar (fractal) time series *x(k,m)* are calculated (int[.] is as an integer function).

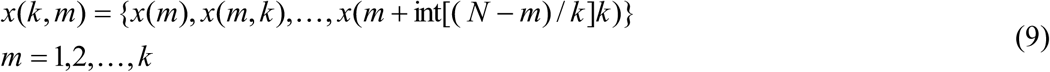
2. We computed the length *L(m,k)* for each of *x(k,m)*.

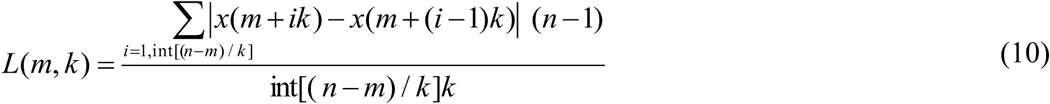
3. *L(m,k)* is then averaged for all *m*. This results in the mean value of the curve length *L(k)*, for each *k*.
4. *Log(L(k))* is plotted versus *log(1/k)*.
5. Fit a straight line to the previous plotted points.
6. The slope of this line is Higuchi’s dimension (HD).

A larger value of Higuchi’s dimension indicates a higher level of complexity.

##### Entropy

The EEG signal *x(t)* is decomposed into brain frequency bands (Delta (0.5-4 Hz), Theta (4.5-8 Hz), Alpha (8.5-16 Hz), Beta (16.5-32 Hz)). The energy of each band is *E*_*1*_, *E*_*2*_, *E*_*3*_, *and E*_*4*_, respectively. *E={E*_*1*_,*E*_*2*_,*E*_*3*_,*E*_*4*_*}* forms an energy distribution in the frequency domain of EEG signals. The corresponding energy entropy (*E*) is calculated as follows [26]:

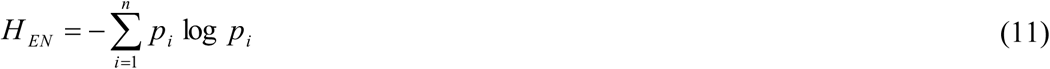

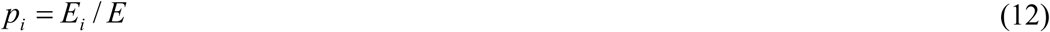

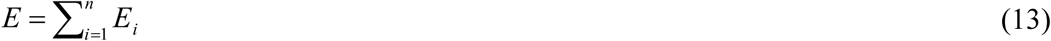

All the mentioned non-linear features were extracted from different frequency bands (i.e., Delta (0.5-4 Hz), Theta (4.5-8 Hz), Alpha (8.5-16 Hz), Beta (16.5-32 Hz)) separately.

##### Lyapunov exponent

The Lyapunov exponent is known as a measure for the detection of the sensitivity of a system to the initial condition. It indicates the average exponential rate of divergence/convergence of the nearby trajectories in the phase space. Positive Lyapunov exponent is associated with systems that are sensitive to the initial condition, for example, chaotic systems. Larger positive exponents reflect a larger time scale in which the dynamics of the system become unpredictable. The largest Lyapunov exponent can be calculated as follows:

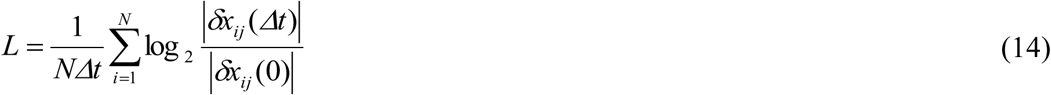

Where, *δx*_*ij*_*(0)=x(t*_*i*_*)-x(t*_*j*_). It is the displacement vector between time points *t*_*i*_ and *t*_*j*_. δ*x*_*ij*_*(Δt)=x(t*_*i*_*+ Δt)-x(t*_*j*_*+ Δt)* is the displacement vector after time *Δt. x(t*_*i*_) and *x(t*_*j*_) are two adjacent vectors in the phase space. As the time increase by *Δt*, two adjacent vectors may diverge. Eq. (14) shows the mean exponential rate of divergence of two initially two adjacent vectors. More details about this index can be found in [31-33].

### 2.6. Classifiers

As mentioned before, one of the goals of this study is to design a system that automatically detects the state of the index finger’s skin with respect to a surface (the contact to, dragging along, and detaching from a surface). To reach to this goal, we examined different classifiers: K-nearest-neighborhood (KNN) (with K=7); linear discriminant analysis (LDA); support vector machine (SVM); and an artificial neural network (ANN) (a feedforward, two hidden layers (with 10 tangent sigmoid neurons in each layers), and linear output neurons).

The inputs of these classifiers were selected features, and their outputs were the mentioned three states. The performance of all classifiers was evaluated using the Leave-One-Out (LOO) cross-validation method.

### 2.7. Statistical analysis

As mentioned before, reducing the number of recording channels and increasing the accuracy of distinguishing between the desired states is important in designing an optimum automatic detection system. Statistical tests were used to identify the features and channels that could best differentiate between desired states. To acquire the significant changes of the features across three states, repeated measure ANOVA with the Geisser-Greenhouse correction and Bonferroni’s multiple comparison test was employed. The significance level was set at 0.05. Features and channels that could make a significant difference between the states were used as the classifier inputs in the automated system design.

The total procedure of the processing stages in designing the system is shown in Fig. 3.

**Fig. 3.**
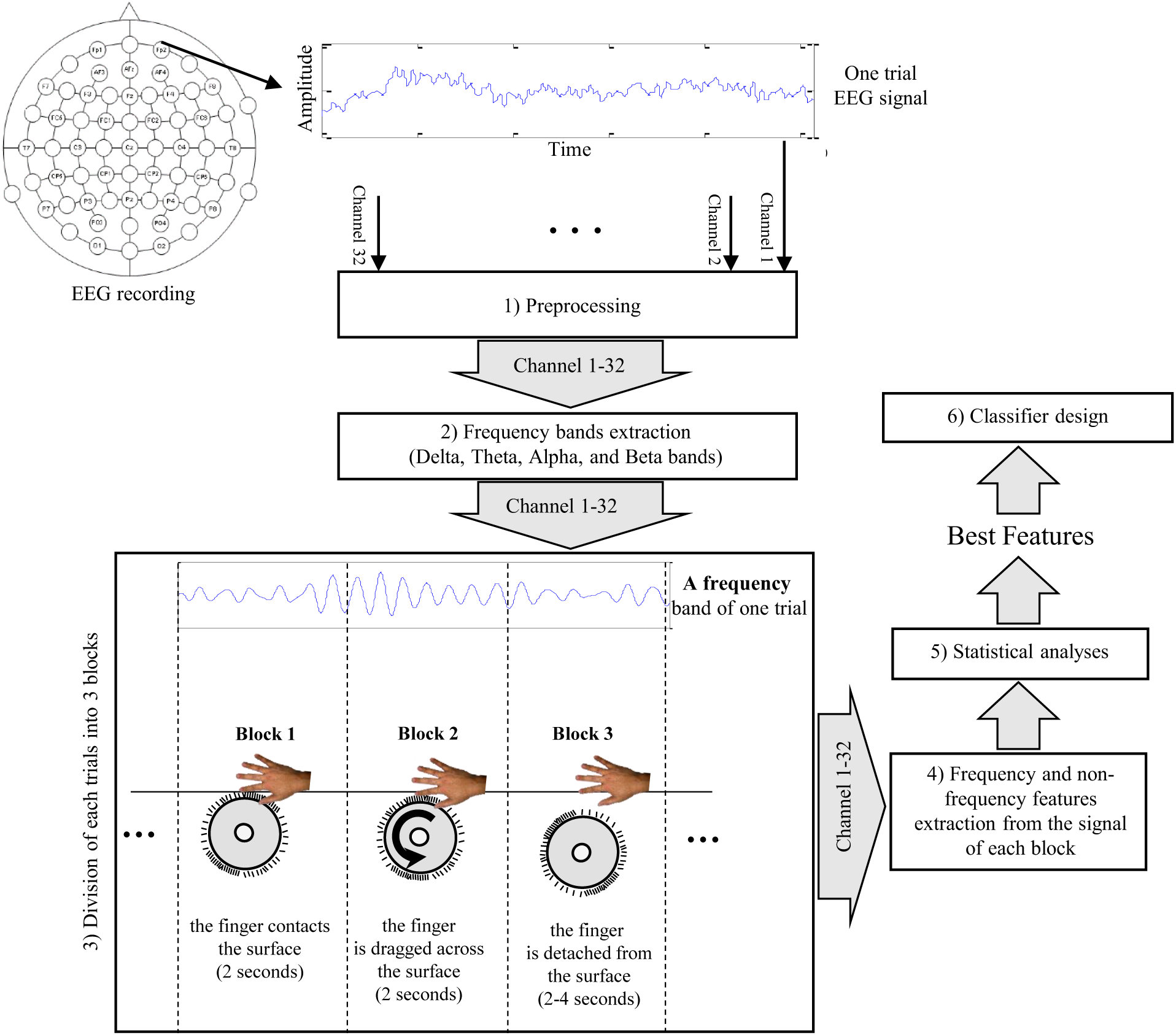
The block diagram of the processing stages in designing the automatic detection system. The preprocessing stage consists of two parts: denoising and artifact rejection from all channels. The second part relates to the extraction of brain frequency bands. In the third stage, the extracted signals of the previous stage are divided into three blocks to explore the effect of dynamic, static, and no friction. This procedure is repeated for both left and right hands. Then, in the fourth stage, different linear and non-linear features are extracted from the signals of the previous stage. The extracted features are investigated by statistical analyses, and features that can distinguish between desired states are selected as best features and are used for design and training the classifier.

## 3. Results

As mentioned before, our aim was to design a system that can automatically detect the states reports in section 2.3 based on the EEG signals:

At first, we investigated the capability of different linear and non-linear features to make a difference between the states statistically. Then, the selected features were used in classifiers. It is worth to be mentioned that the statistical analysis for each surface (rough, semi-rough, and soft) and for left and right hands have been performed separately. The best features in all examined surfaces in both hands were given as inputs to the classifier.

### 3.1. Linear features

The results of the Bonferroni’s multiple comparison tests after the repeated measure ANOVA with the Geisser-Greenhouse correction showed that the linear features could not distinguish between the three desired states (i.e., Blocks 1, 2, and 3) in both hands. Therefore, it seems that the use of linear features for the design of the desired automatic detection system is not an appropriate decision. At the next stage, we examined the non-linear features that were extracted from each brain wave signal. The results are reported in the next section.

### 3.2. Non-linear features

As mentioned before, we extracted Hurst exponent, Higuchi’s dimension, Lyapunov exponent, and entropy from the decomposed EEG signals of each 32 recording channel. To visualize the results and easier selection of best features and channels, we have shown the results of the statistical analyses in the following figures. Fig. 4 shows features and brain channels that are most capable of detecting the expected states (Block1 (B1), Block2 (B2), and Block3 (B3)) from each other in touching the rough (R) surface with left (L) hand (R-L).

**Fig. 4.**
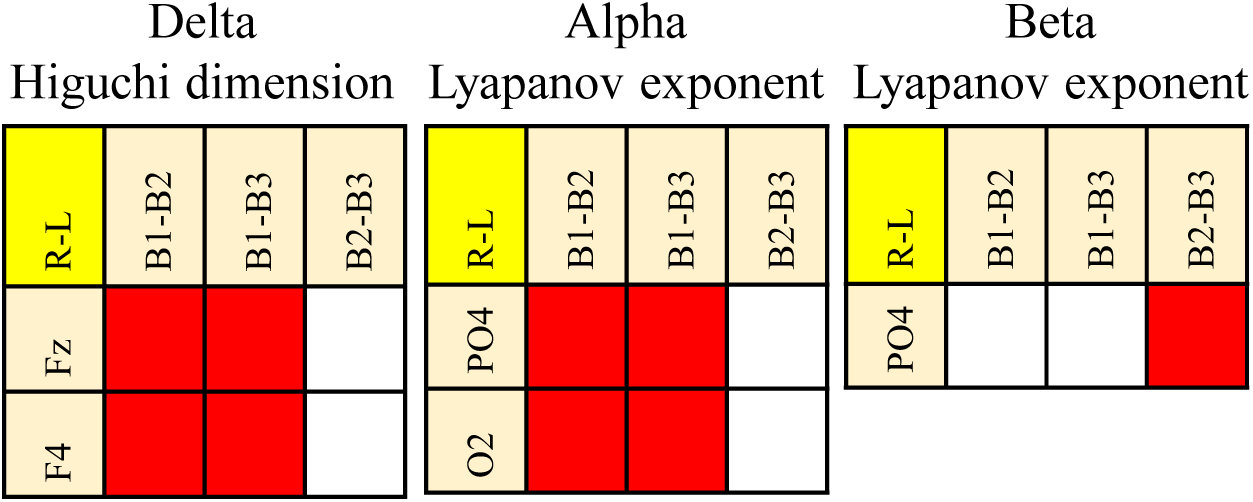
Features and brain channels that are most capable of detecting the expected states (Block1 (B1), Block2 (B2), and Block3 (B3)) from each other in touching the rough (R) surface with left (L) hand (R-L).

According to Fig. 4, no feature can separate three states while touching the rough surface with the left hand. However, Alpha band Lyapunov exponent of the PO4 channel can differentiate between Block1 (B1) and Block2 (B2), and can also detect Block1 (B1) from Block3 (B3). On the other hand, the Beta band Lyapunov exponent of the PO4 channel can be used to distinguish Block2 (B2) from Block3 (B3). Therefore, the simultaneous use of the calculated Alpha and Beta bands Lyapunov exponent of the PO4 channel can distinguish between all three states. These features were used as one of the inputs of the classifier in the designed system. Fig. 5 shows the value of these features in the three blocks of each trial.

**Fig. 5.**
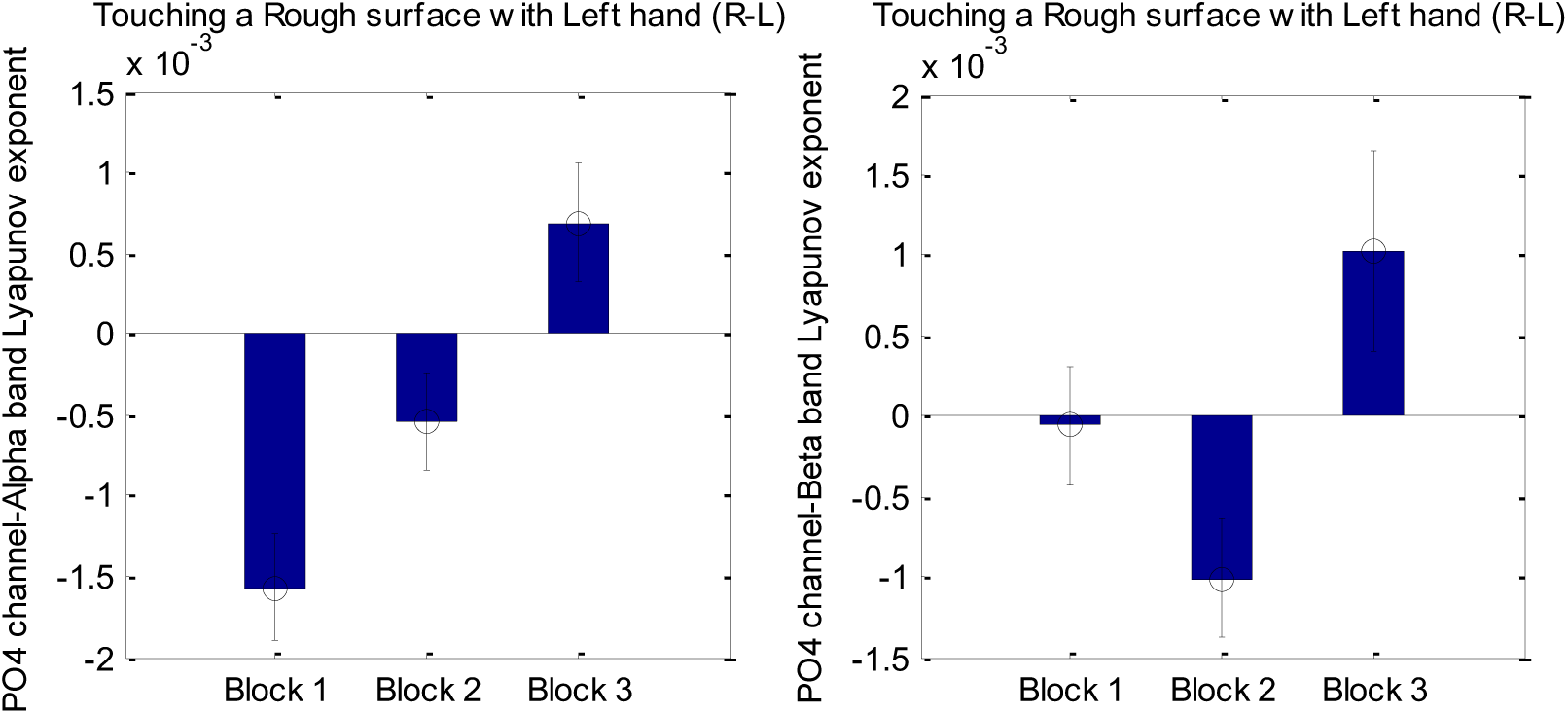
The value of Alpha (right panel) and Beta (left panel) bands’ Lyapunov exponent calculated from the PO4 channel’s signal in touching a rough surface with the left hand (R-L).

According to Fig. 5, it seems that the value of the Alpha and Beta bands’ Lyapunov exponent in blocks 1 and 2 is negative, and in block 3 is positive. This result suggests some information about the dynamics of the tactile system that will be discussed in the discussion part.

Fig. 6 shows features and brain channels that are most capable of detecting the expected states (Block1 (B1), Block2 (B2), and Block3 (B3)) from each other in touching the semi-rough (SR) surface with left (L) hand (SR-L).

**Fig. 6.**
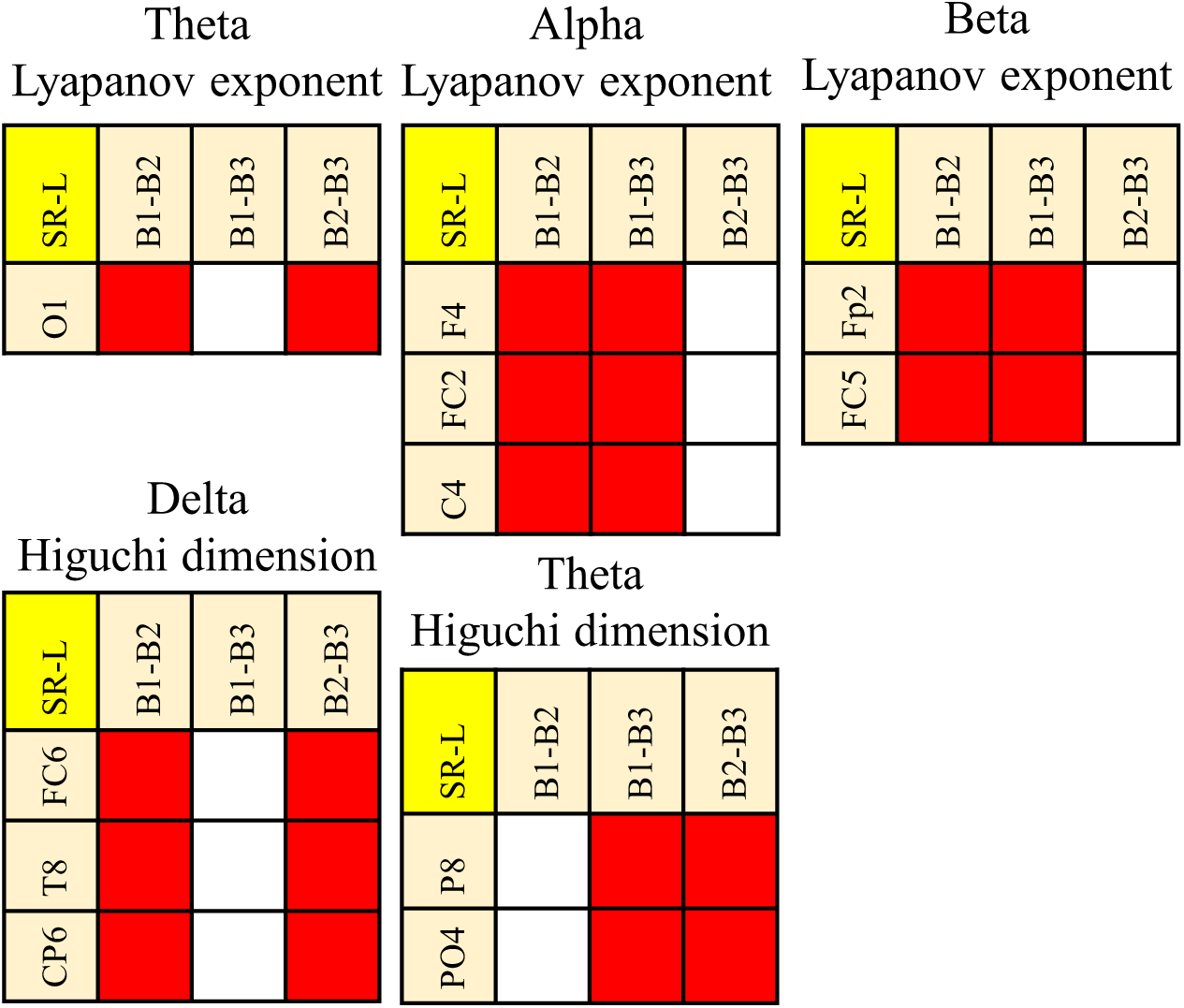
Features and brain channels that are most capable of detecting the expected states (Block1 (B1), Block2 (B2), and Block3 (B3)) from each other in touching the semi-rough (SR) surface with left (L) hand (SR-L).

Like the previous mode (R-L), no feature can differentiate between all three blocks. However, the simultaneous use of the T8 Delta band’s Higuchi’s dimension and PO4 Theta band’s Higuchi’s dimension can distinguish the blocks. According to this figure, in addition to these features, the simultaneous use of other features can also be used. However, to reduce the numbers of recording channels, we tried to select features that have common channels with ones that have been selected for other modes (R-L, SR-L, S-L, R-R, SR-R, and S-R). Fig. 7 shows the value of the T8 Delta band’s Higuchi’s dimension and PO4 Theta band’s Higuchi’s dimension in the three blocks of each trial.

**Fig. 7.**
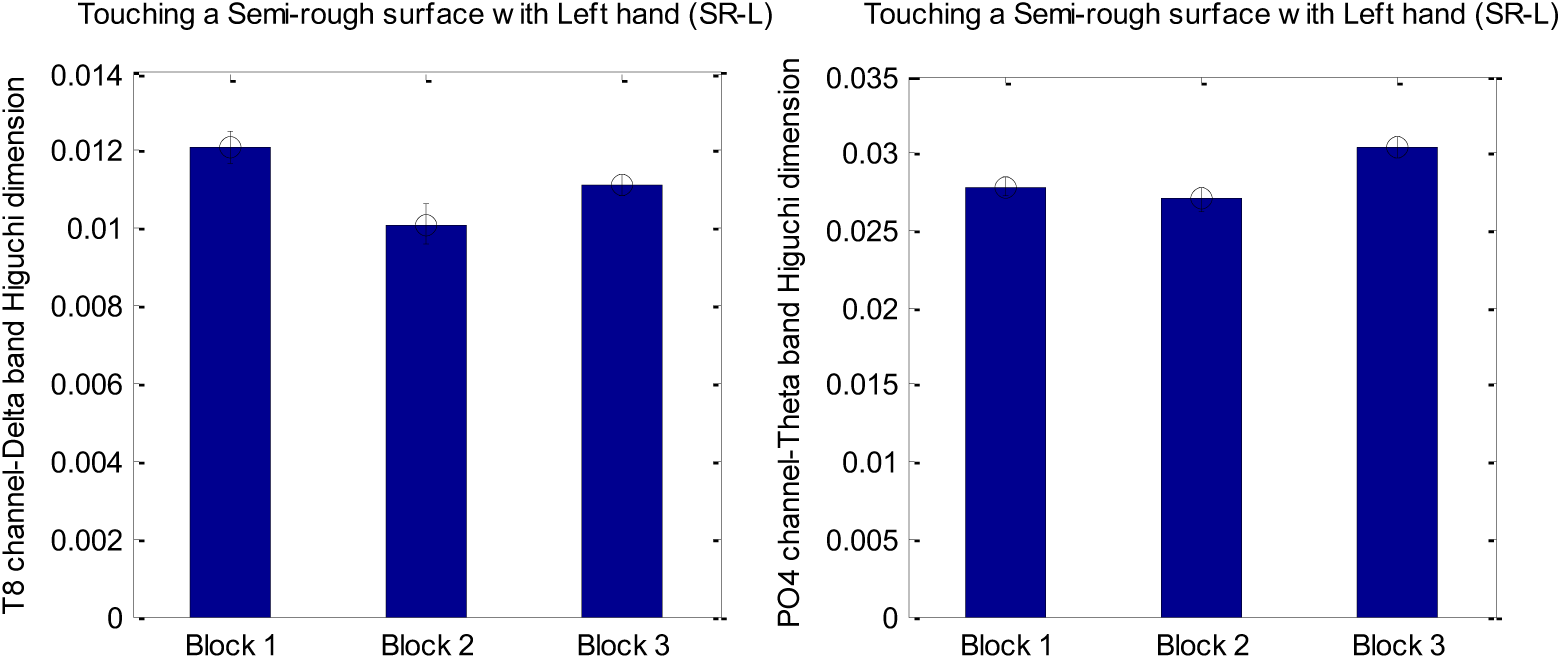
The value of the T8 channel Delta band’s Higuchi’s dimension (left panel) and PO4 channel Theta band’s Higuchi’s dimension (right panel) in touching a semi-rough surface with the left hand (SR-L).

Fig. 8 shows Features and brain channels that are most capable of detecting the expected states (Block1 (B1), Block2 (B2), and Block3 (B3)) from each other in touching the soft (S) surface with left (L) hand (S-L).

**Fig. 8.**
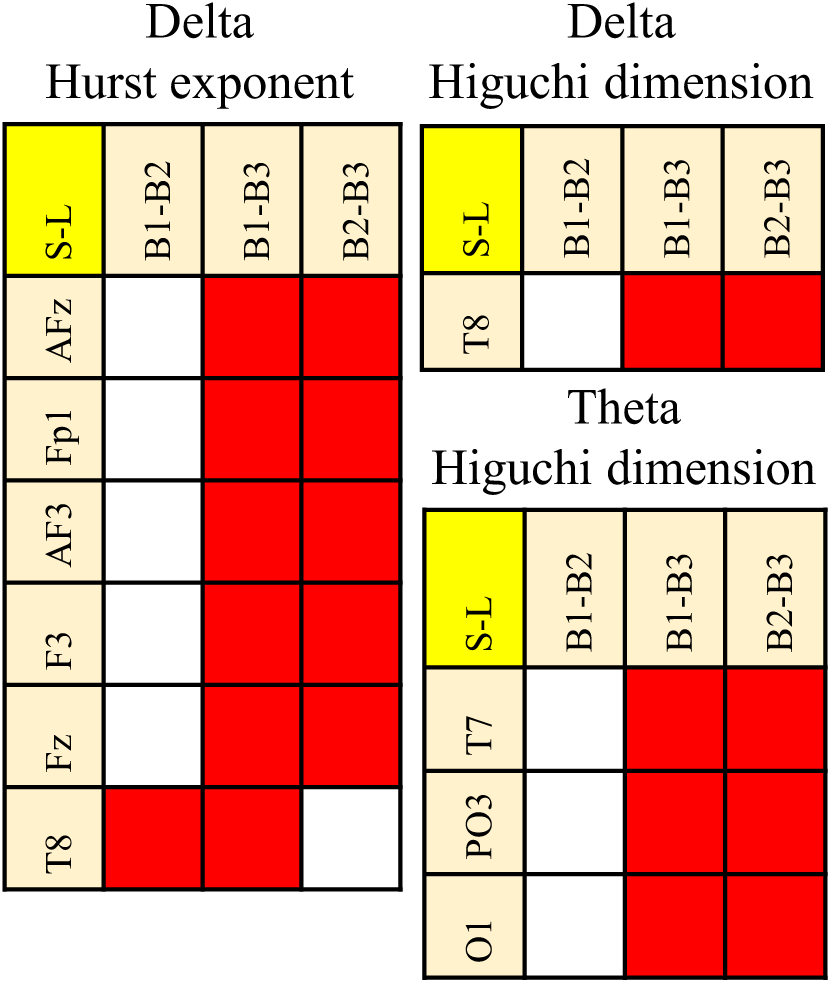
Features and brain channels that are most capable of detecting the expected states (Block1 (B1), Block2 (B2), and Block3 (B3)) from each other in touching the soft (S) surface with left (L) hand (S-L).

According to Fig. 8, the simultaneous use of the value of Delta bands’ Hurst exponent calculated from Fz and T8 channels can distinguish between three blocks of the test. One of the other possible options is the simultaneous use of the value of Delta bands’ Hurst exponent and Higuchi’s dimension calculated from the T8 channel. To reduce the number of recording EEG channels, we selected the second option. The features in the second option were selected as inputs of the classifier. Fig. 9 shows the mean value of these features in three blocks.

**Fig. 9.**
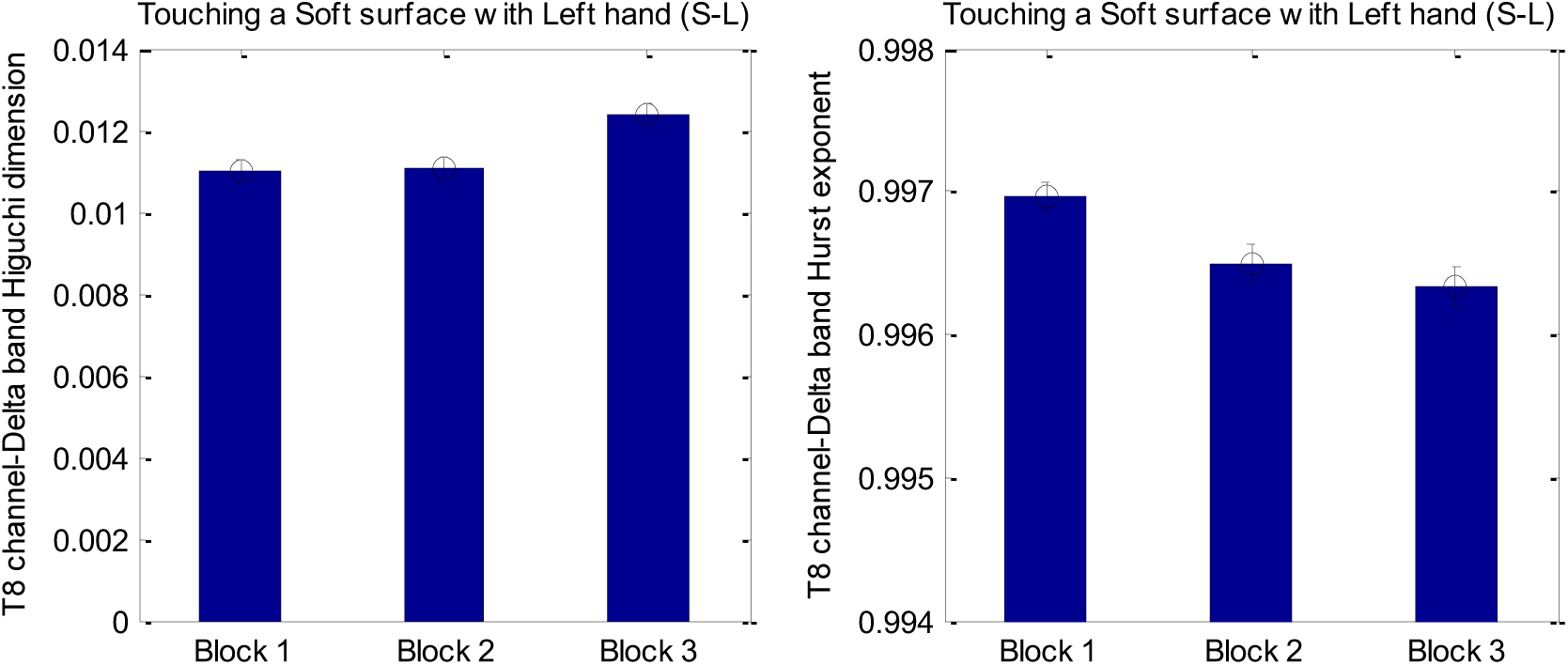
The value of Delta bands’ Hurst exponent (right panel) and Higuchi’s dimension (left panel) calculated from the T8 channel’s signal in touching a soft surface with the left hand (S-L).

Fig. 10 shows features and brain channels that are most capable of detecting the expected states (Block1 (B1), Block2 (B2), and Block3 (B3)) from each other in touching the rough (R) surface with right (R) hand (R-R).

**Fig. 10.**
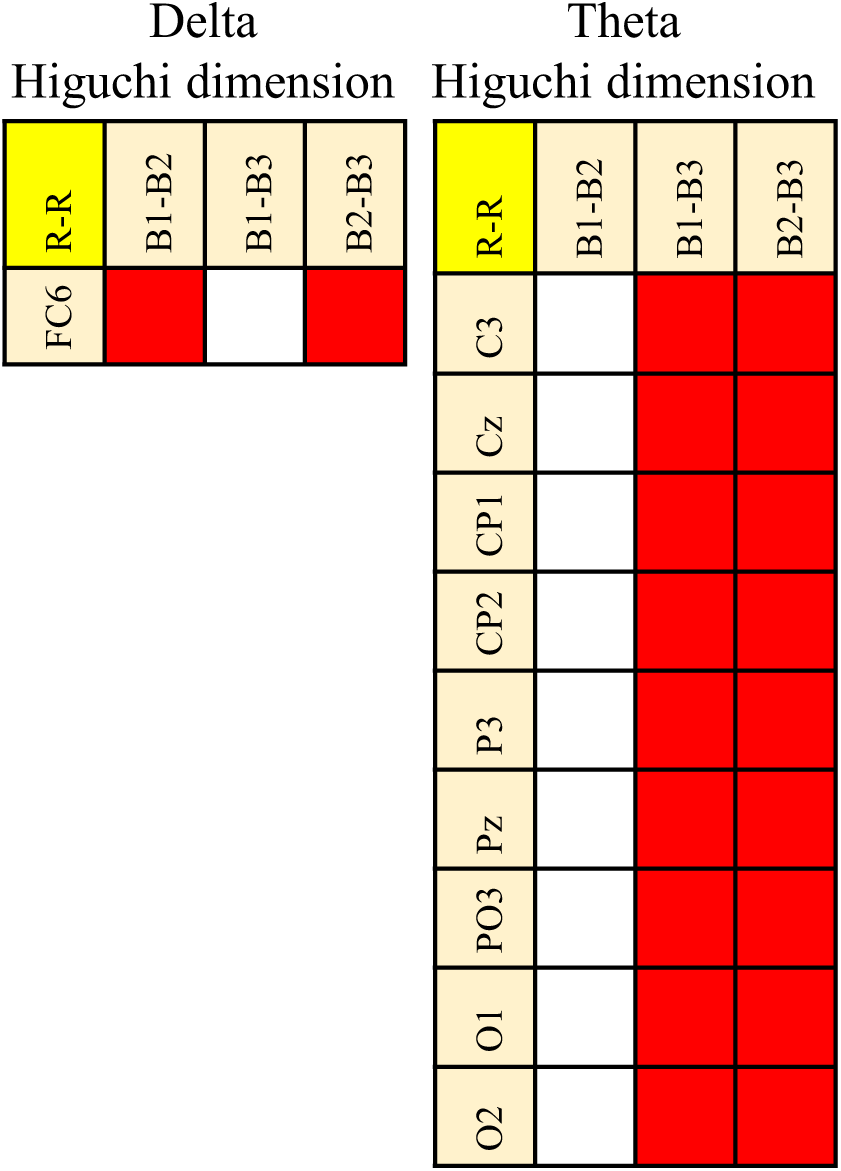
Features and brain channels that are most capable of detecting the expected states (Block1 (B1), Block2 (B2), and Block3 (B3)) from each other in touching the rough (R) surface with right (R) hand (R-R).

According to Fig. 10, the simultaneous use of the FC6 channel Delta band’s Higuchi’s dimension and O1 channel Theta band’s Higuchi’s dimension can be used to differentiate among the blocks of each trial in touching the rough (R) surface with right (R) hand (R-R). Fig. 11 shows the mean value of these features in three blocks.

**Fig. 11.**
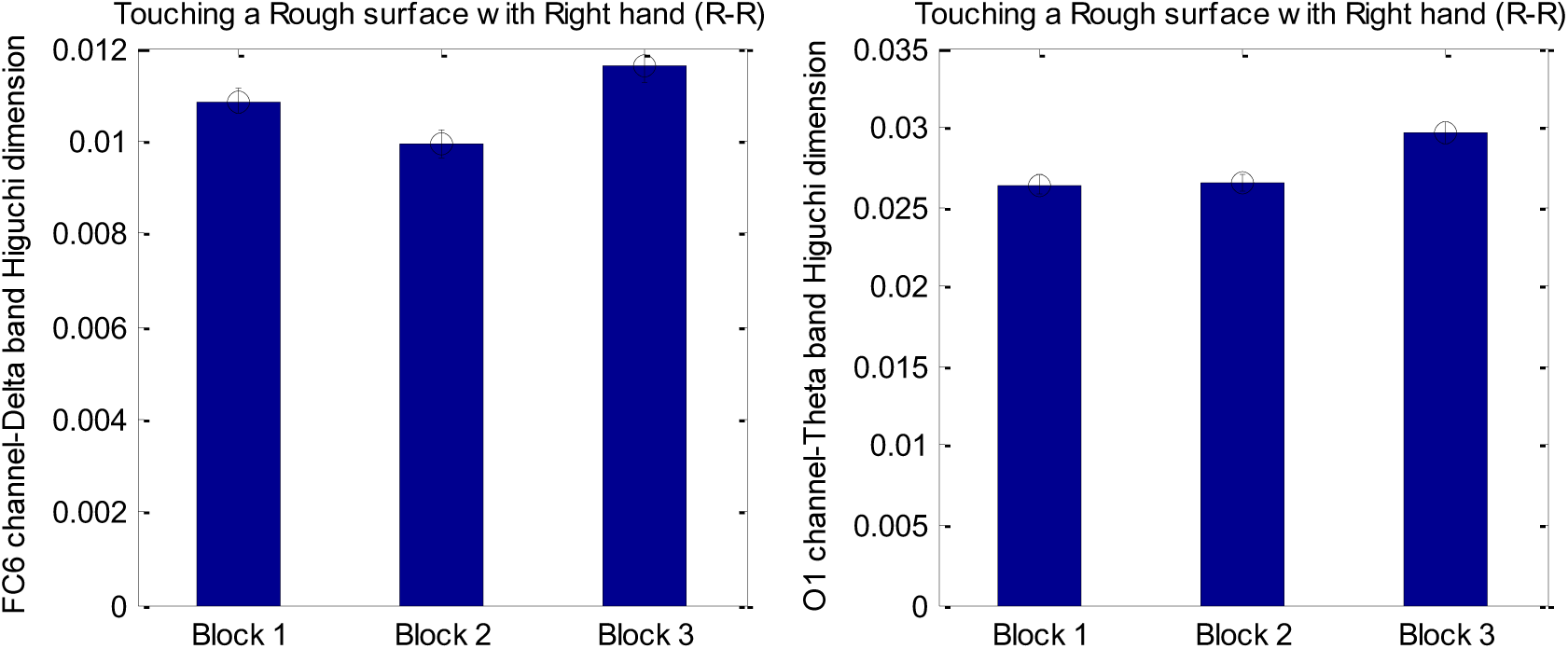
The value of the FC6 channel Delta bands’ Higuchi’s dimension (left panel) and O1-channel Theta band’s Higuchi’s dimension (right panel) in touching a rough surface with the right hand (R-R).

The reason for the selection of O1, among other possible channels, is that this channel has also been used in the next mode (SR-R).

Fig. 12 shows Features and brain channels that are most capable of detecting the expected states (Block1 (B1), Block2 (B2), and Block3 (B3)) from each other in touching the semi-rough (SR) surface with right (R) hand (SR-R).

**Fig. 12.**
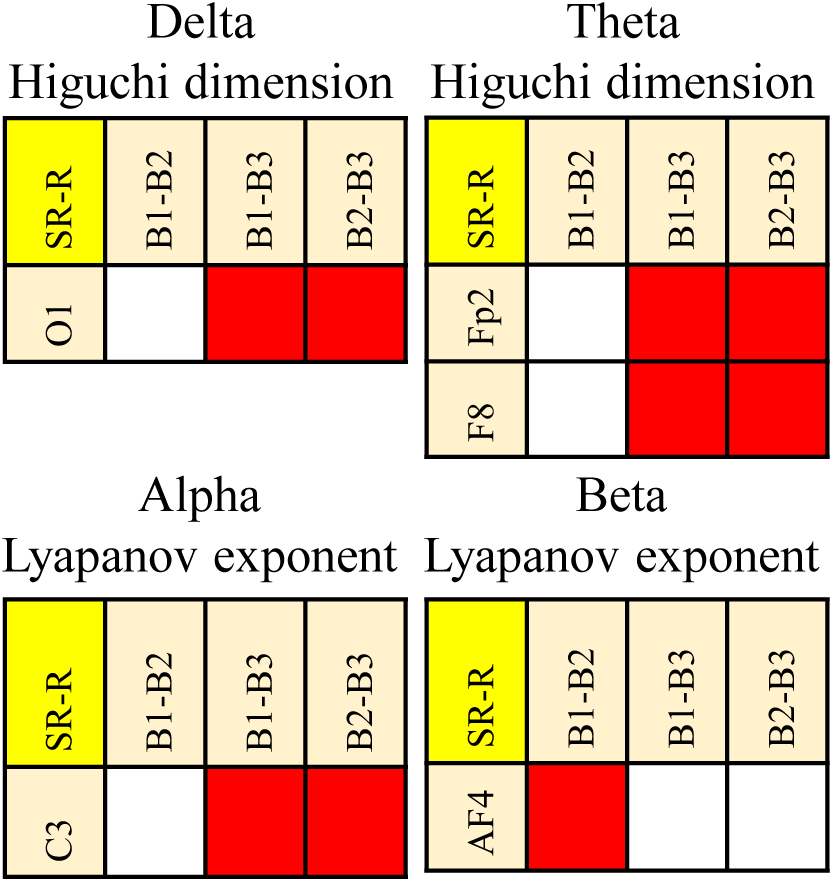
Features and brain channels that are most capable of detecting the expected states (Block1 (B1), Block2 (B2), and Block3 (B3)) from each other in touching the semi-rough (SR) surface with right (R) hand (SR-R).

According to Fig. 12, the simultaneous use of the AF4 channel Beta band’s Lyapunov exponent and the O1 channel Delta band’s Higuchi’s dimension can distinguish three blocks of the trials. The reason for the selection of O1 among other channels that can differentiate between block 3 and other blocks is that O1 has also been selected in previous mode (R-R). The selection of common channels in different modes leads to the decrement of recording channels. Fig. 13 shows the mean value of these features in three blocks.

**Fig. 13.**
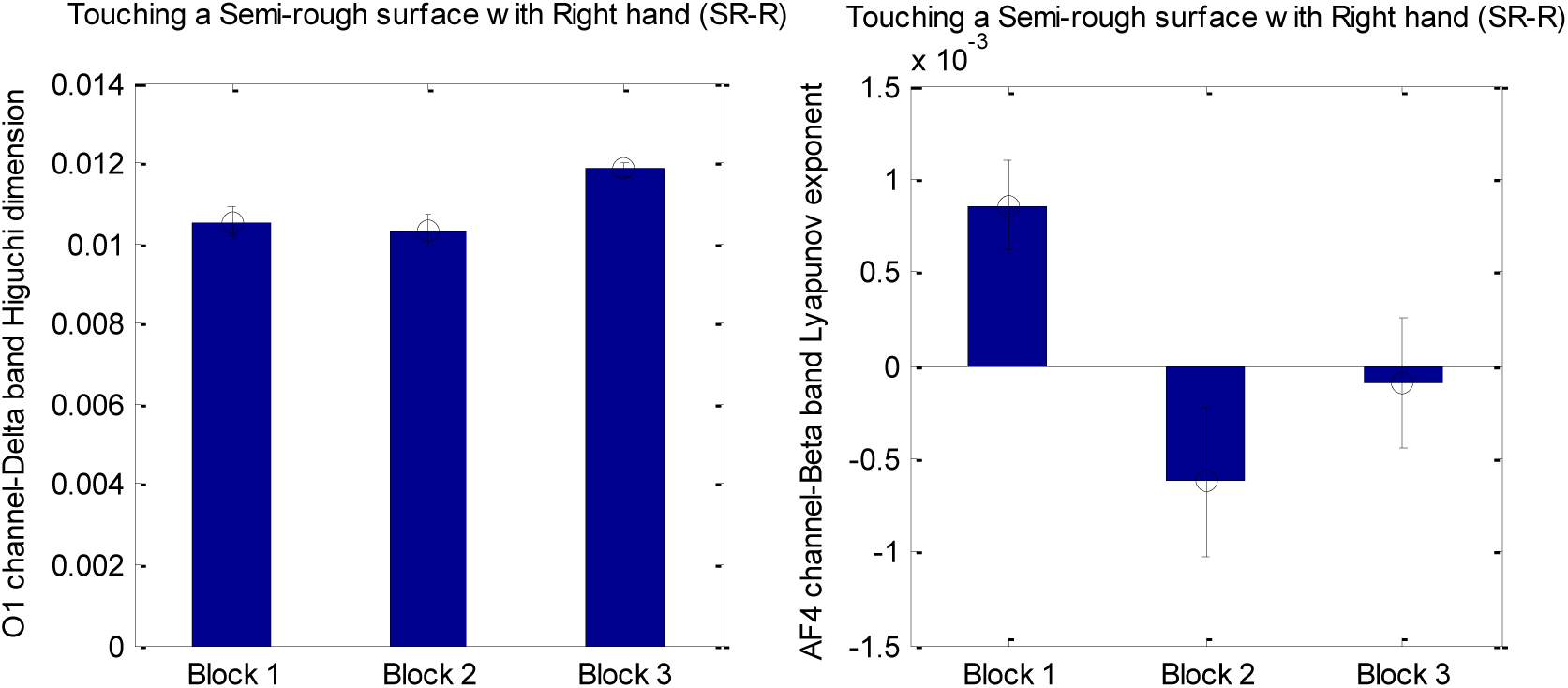
The value of the O1 channel Delta bands’ Higuchi’s dimension (left panel) and AF4-channel Beta band’s Lyapunov exponent (right panel) in touching a semi-rough surface with the right hand (SR-R).

Fig. 14 shows Features and brain channels that are most capable of detecting the expected states (Block1 (B1), Block2 (B2), and Block3 (B3)) from each other in touching the soft (S) surface with right (R) hand (S-R).

**Fig. 14.**
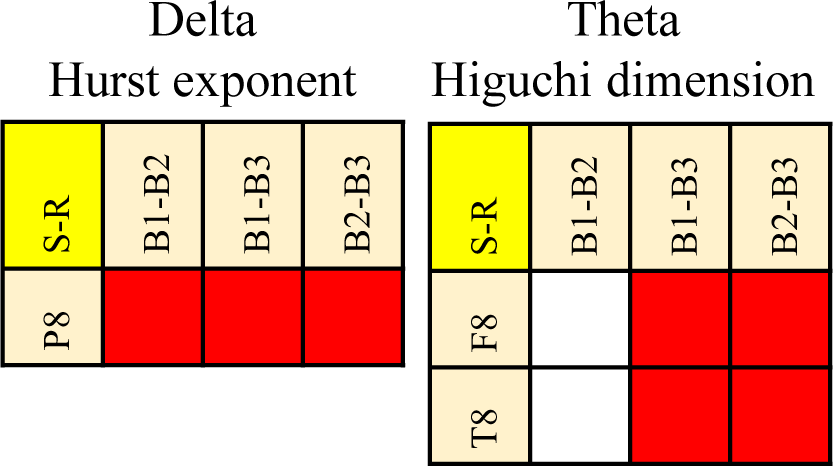
Features and brain channels that are most capable of detecting the expected states (Block1 (B1), Block2 (B2), and Block3 (B3)) from each other in touching the soft (S) surface with right (R) hand (S-R).

Considering Fig. 14, it seems that the value of the Delta band’s Hurst exponent recorded from channel P8 can differentiate the blocks. Therefore, this feature was also used as one of the inputs of the classifier. Fig. 15 shows the mean value of this feature in three blocks.

**Fig. 15.**
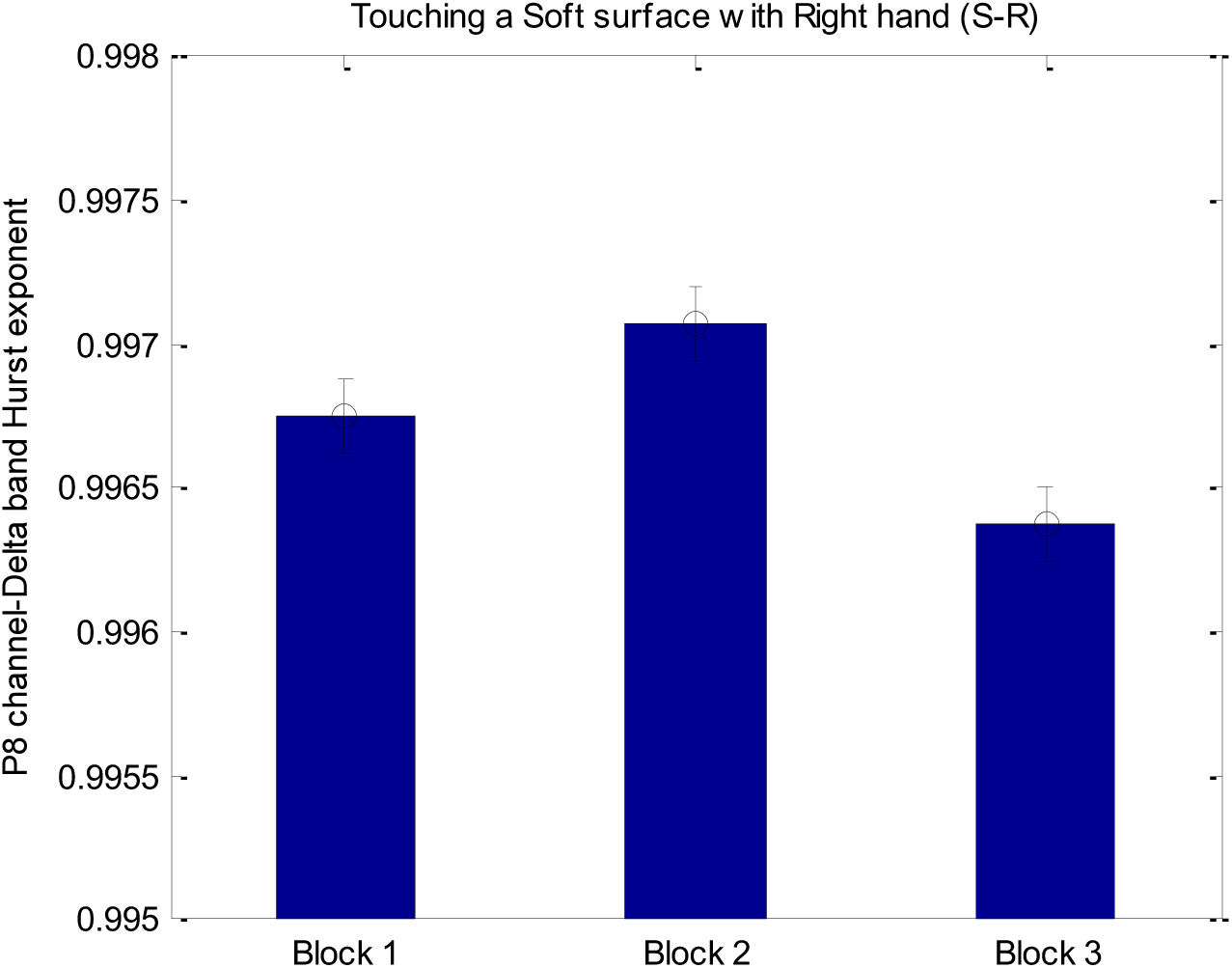
The value of the P8 channel Delta band’s Hurst exponent in touching a soft surface with the right hand (S-R).

### 3.3. Classifier

According to the results of the statistical analyses described in the previous section, we have chosen the following best features.

1. PO4 Alpha band’s Lyapunov exponent
2. PO4 Beta band’s Lyapunov exponent
3. T8 Delta band’s Higuchi’s dimension
4. PO4 Theta band’s Higuchi’s dimension
5. T8 Delta band’s Hurst exponent
6. FC6 Delta band’s Higuchi’s dimension
7. O1 Theta band’s Higuchi’s dimension
8. O1 Delta band’s Higuchi’s dimension
9. AF4 Beta band’s Lyapunov exponent
10. P8 Delta band’s Hurst exponent

These are the features and channels that have been able to make the most statistically significant difference between the three states concerning the other features and channels. As shown in Fig. 16, the selected channels mostly located on the right side of the brain.

**Fig. 16.**
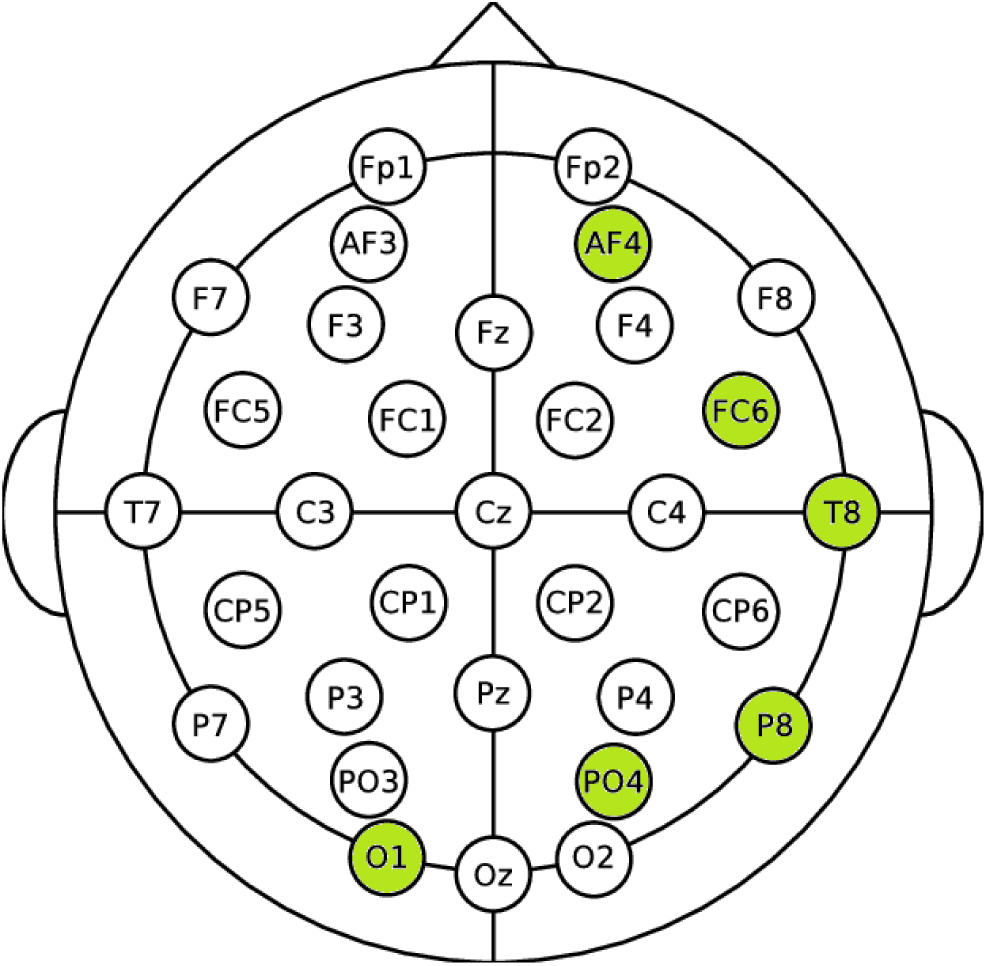
A schematic of selected brain channel distribution

These features were given as inputs to the classifiers. The outputs of the classifiers are the three states (Blocks 1, 2, and 3) mentioned in section 3.2. That is, the classifier, which is one of the main parts of our automatic detection system, takes the above EEG features as inputs and automatically detect which block (contacting, dragging, or detaching) of the test has occurred. Table 1 shows the performance of different classifiers.

**Table 1.** Percentage error of each classifier in detecting each block. The errors relate to the test data based on Leave-One-Out (LOO) cross-validation method.

|  | KNN | LDA | SVM | ANN |
| --- | --- | --- | --- | --- |
| Block 1 | 44% | 37% | 29% | 40% |
| Block 2 | 43% | 28% | 59% | 34% |
| Block 3 | 48% | 37% | 57% | 33% |
| total | 45% | 34% | 49% | 66% |

According to this table, it seems that LDA performance is better than others are. In addition, SVM has had fewer errors in the detection of Block 1, but the total performance of LDA was better than others. Fig. 17 shows the distribution of the ten selected features in three blocks of the experiment.

**Fig. 17.**
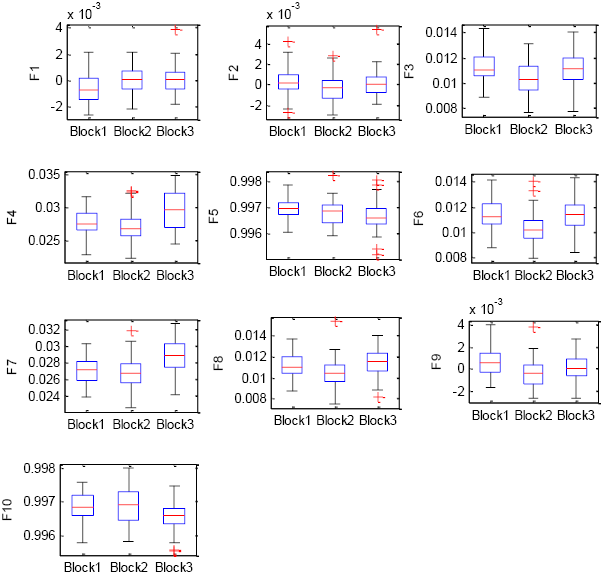
Boxplot diagrams of selected features (F1-F10) in three Blocks of the experiment

Although the selected ten features were statistically different between some of the experiment’s blocks, as shown in Fig. 17, the overlap between the values of each feature among the blocks caused the classification error.

To decrease the overlap and increase the accuracy of classification, we calculated two linear combinations of the selected features that made the most significant difference between the three blocks. The coefficient of each feature in these linear combinations is the first two vectors of coefficients for the canonical variables in one-way multivariate analysis of variance, and they are scaled so that the within-group variance of the canonical variables is one. Using these two features, the classifiers were trained again. Table 2 shows the performance of different classifiers based on these two features.

**Table 2.** Percentage error of each classifier in detecting each block. The errors relate to the test data based on the Leave-One-Out (LOO) cross-validation method according to the new features.

|  | **KNN** | **LDA** | **SVM** | **ANN** |
| --- | --- | --- | --- | --- |
| **Block 1** | 26% | 37% | 29% | 70% |
| **Block 2** | 24% | 22% | 50% | 74% |
| **Block 3** | 42% | 33% | 37% | 59% |
| **total** | 31% | 31% | 39% | 68% |

According to the results reported in Table 2, the performance of the classifiers has improved slightly, and the LDA classifier still performs the best. Fig. 18 shows the distribution of test data (blue circles) and the output of different classifiers (red stars).

**Fig. 18.**
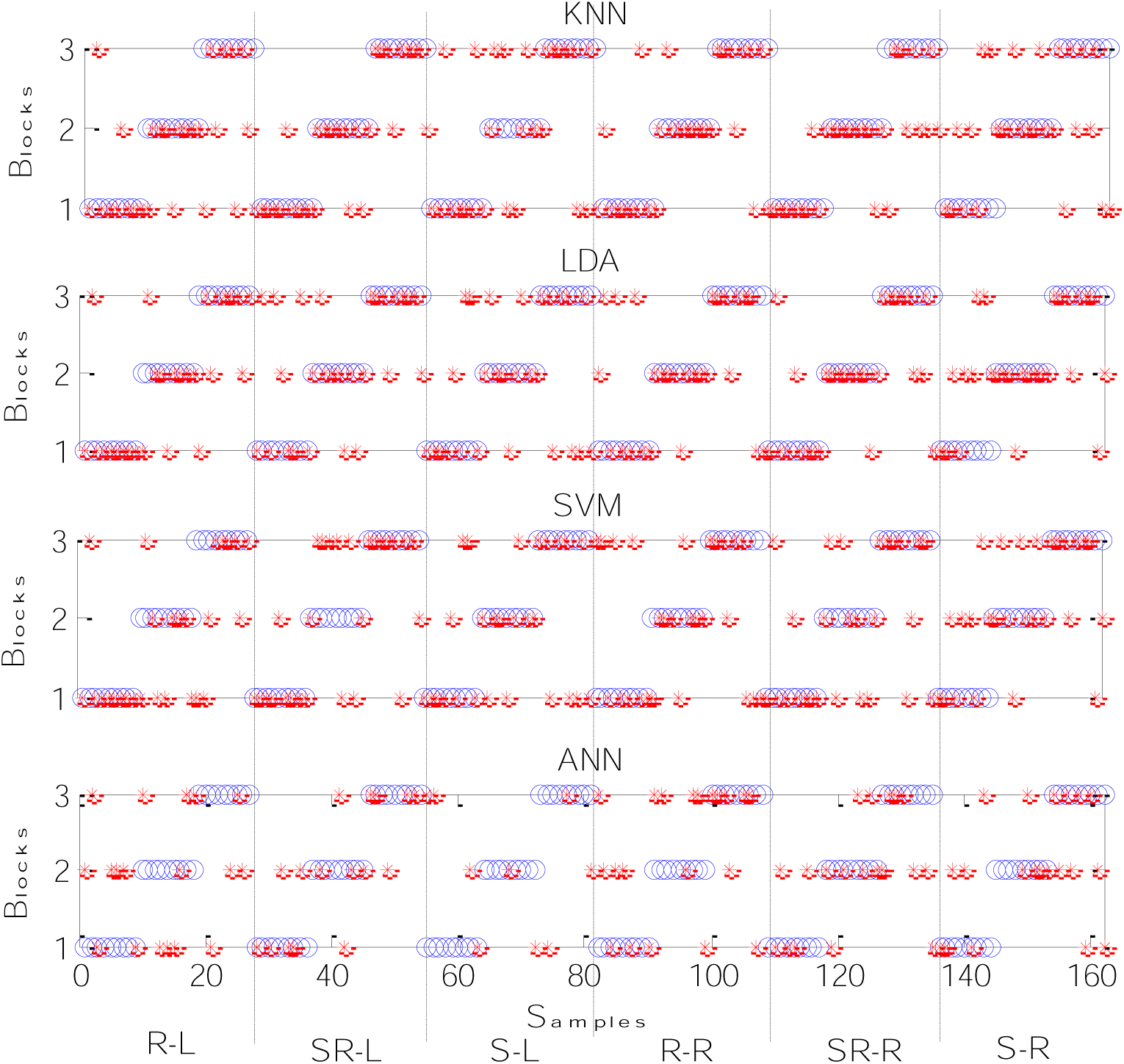
Distribution of test data (blue circles) and the output of different classifiers (red stars); R-L: rough surface touched with left hand; SR-L: semi-rough surface touched with left hand; soft surface touched with left hand; rough surface touched with rough hand; SR-L: semi-rough surface touched with rough hand; soft surface touched with rough hand.

Fig. 18 indicates that the errors of classifiers in the detection of each block do not relate to a special roughness level or handedness because the distribution of incorrectly detected samples (stars) in all cases was almost the same.

## 5. DISCUSSION

In this study, the main goal was to design a system for automatic detection of the creation of friction between the index finger skin and a surface. The experiment consisted of three blocks so that in each block, the position of the fingertip changed relative to the surfaces. In the first block, the fingertip was in direct contact with the surface without movement (static friction). In the second block, the surface was pulled out underneath the fingertip (dynamic friction). Finally, in the third block, the surface was detached from the fingertip (no friction). We hypothesized that switching between these blocks may affect the brain dynamics. Considering the other studies, the variation of the dynamics of the neural systems and their responses to tactile friction has been reported in different conditions. However, to the best of our knowledge, there is no study to explore the variations of the brain’s electrical responses under the alternation of the friction type (i.e., block1: static friction, block2: dynamic friction, and block3: no friction) during the experiment. Moreover, in tactile studies, no system has been suggested to detect friction between the skin and a surface automatically. In addition, we studied the effect of tactile friction on non-linear features of EEG signals for the first time. Results of the current study showed that switching between the states (static friction (Block 1), dynamic friction (Block 2), and no friction (Block 3)) had no significant effects on the linear features of EEG signals (the energy of different brain waves). However, the changes of EEG linear features, especially Theta and Alpha band energy, have been reported in tactile studies related to the roughness detection [11-14].

Theta and Alpha bands have been associated with our focus on internal signals [34, 35]. Therefore, it can be suggested that tactile friction sensation or perception does not affect brain resources that are associated with the focus on internal signals. However, some of the non-linear features extracted from different brain waves were changed in three blocks of the experiment.

As shown in Fig. 5, the value of the Alpha and Beta bands’ Lyapunov exponent in blocks 1 and 2 is negative, and block 3 is positive. Positive values of Lyapunov exponent indicate that the signal may be chaotic, and its negative values indicate that the signal is not chaotic. Blocks 1 and 2 are respectively associated with contacting and dragging the finger on the surface. Block 3 relates to the detaching the finger from the surface. The obtained negative values for the first and second blocks that engaged the tactile system can suggest the existence of non-chaotic dynamics and predictability characteristics in EEG signals recording during the tactile friction sensation. This result is consistent with the results obtained from Hurst’s exponent analysis, which reveals the existence of dependency in the signals.

Fig. 9 and Fig. 15 show that Hurst exponent values in all states are close to one. These values show the existence of long-range dependency in tactile EEG signals.

Considering the values of Higuchi’s dimension in Figs Fig. **8**, Fig. **11**, and Fig. **13** indicates that the value of Higuchi’s dimension in Theta band is higher than that of in Delta band. Higuchi’s dimension is an index of the complexity of the system. Therefore, it seems that in Theta band, the complexity of the tactile system is higher than that of in Delta band. Results also indicated that tactile friction sensation does not affect the entropy of the EEG signals.

These selected non-linear features were given as input to different classifiers to design a system that automatically could detect the desired states (static, dynamic, and no tactile friction). Using LDA classifier and the calculation of 10 features from the signals of six EEG channels (O1, PO4, P8, T8, FC6, and AF4) we can automatically detect when there is a static, dynamic, and no tactile friction between the skin and surfaces with different levels of roughness with error rate of 31%. Although this is a preliminary result and it is required to be investigated further with larger data sets, the results can open new ways in the development of novel tactile BCI systems for future works such as follows:

- The knowledge about the effect of friction type on brain dynamics can be used in developing robotic hands in disabled people. It is tried to develop these artificial hands in a way that leads to a real feeling of the occurrence of friction between the skin and surfaces. That is, using brain stimulation methods, it is tried to change the brain dynamics so that an artificial sense of friction is induced in the brain. Moreover, the individuals who wear such a robotic hand can detect the contact of his/her hand with different surfaces.
- In virtual reality games or BBI systems, the proposed automatic detection system can also be used to induce a sense of tactile friction. That is, the designed automatic system detects the state of the friction between an individual’s skin and a surface. Then, using a computer-brain interface (CBI), the brain dynamics of other participants are changed so that they virtually sense the same state of friction that the first individual feels.
- Another application of these tactile BCI systems can be in the textile industry and neuro-economic fields. An important point in industrial design and manufacturing is pleasant touching. It has been reported that there is a correlation between tactile friction and the touch comfort level [36]. The results of the current study showed that the type of friction also affected brain dynamics and, consequently, the tactile friction perception. It can be concluded that considering the changes of EEG signals, the first contact of the skin with an object is as important as moving the skin on that object. Therefore, the study of the brain response to the first contact of the skin with a surface may also help scientists in textile industry to produce more comfortable clothing and neuro-economic fields to form a pleasure sense in customers in the first contact of their skin with the products and keeping this feeling while the customer moves his\her finger on the product.

## Declarations of interest

none

